# Big-team science reveals that patterns of inhibitory control variation in teleost fishes differ from those in mammals and birds

**DOI:** 10.64898/2026.09.22.753528

**Authors:** Laurent Prétôt, Elia Gatto, Tyrone Lucon-Xiccato, Samina Abedi, Christian Agrillo, Pablo Arechavala-López, Maria P. Arteaga-Avendaño, Leonardo Ávila, Alexis Berry, Benjamin C. Bluck, Culum Brown, Redouan Bshary, María J. Cabrera-Álvarez, Gabriella Dennis, Vanessa Guerrini, Yseult Héjja-Brichard, Kyndal B. Irwin, Nicola Jung, Morgan Leeper, Tamra C. Mendelson, Kasturi Mondal, Inês Neves, Cait Newport, Alessandra Pecunioso, Shane Rance, João L. Saraiva, Ronen Segev, Levi Storks, Justin Yeager, Zegni Triki

## Abstract

Understanding the evolution of cognition is fundamental to understanding the origins and nature of our own cognitive abilities, with cross-species comparisons providing a cornerstone of this endeavor. Yet, comparative cognition so far has largely focused on endothermic vertebrates, particularly mammals and birds, while ectotherms—representing a substantial proportion of vertebrate diversity—remain relatively understudied. Moreover, variation in experimental methods makes existing data difficult to compare across species. Big-team science offers a way to overcome these limitations by enabling standardized cognitive testing across broad taxonomic scales. Here, we apply this approach to fishes, using inhibitory control as a proof of concept for standardized comparative testing in 444 individuals across 22 teleost species spanning 19 genera. We demonstrate that standardized cognitive testing across such diversity is feasible, while also revealing substantial heterogeneity in the sample. Performance varied markedly among species, but, strikingly, showed only a weak phylogenetic signal, contrasting with patterns previously reported in mammals and birds. In addition, we detected great within-species variation, including differences across research groups, suggesting that performance may reflect not only species-level evolutionary history but also ontogenetic experience and the capacity to adjust decision rules to local conditions. Together, our findings highlight the importance of both evolutionary and environmental sources of cognitive variation and demonstrate that extending standardized comparative approaches beyond traditionally studied endotherms can reveal a more dynamic and complex picture of cognitive evolution across the vertebrate lineage.

## INTRODUCTION

Animals rely on cognition to acquire and process information about their physical and social environments and use it to make behavioral decisions that meet their ecological needs. Understanding how and why these abilities vary across species is therefore central to explaining the evolution of cognition: which selective pressures favor particular cognitive abilities, and how species adjust to the specific challenges they face. Traditionally, comparative psychology has addressed these questions by comparing cognitive performance across independent studies and examining small, taxonomically narrow samples. More recently, evidence has emerged that even subtle methodological differences in task implementation can significantly affect assessments of cognitive abilities (e.g., [1–3]; see [4–9]). This recognition has fueled the rise of a new framework called “big-team science”, which consists of large-scale, coordinated comparative projects that use standardized paradigms within and across taxa (e.g., [10, 11]).

Big-team science has opened new opportunities for comparative cognition by coordinating standardized experiments across species, enabling variation in cognitive performance to be examined at a taxonomic scale that is difficult to achieve within individual research groups. For example, a large-scale comparison of short-term memory across primates previously identified a strong phylogenetic signal, with closely related species showing more similar cognitive performance [12, 13]. This is in contrast with earlier studies showing that behavioral traits often exhibit weaker phylogenetic signals than morphological traits [14].

Another large-scale comparative study—highly relevant to the current work—investigated inhibitory control across 36 species of mammals and birds (including 23 primates and 7 birds) using two paradigms, cylinder and A-not-B tasks [15]. The main finding was a strong phylogenetic signal, with closely related species performing more similarly than distantly related species. Additionally, absolute brain size was a key predictor of performance across all species in the study. Interestingly, inhibitory control within primates was also associated with dietary breadth but not group size; however, other work has linked variation in inhibitory control performance to social systems rather than phylogenetic relationships and feeding ecology [16]. Taken together, these findings suggest that the evolutionary drivers of cognitive variation likely differ across both taxonomic groups and cognitive domains.

Despite the insights gained from large-scale comparative studies, our understanding of cognitive evolution remains heavily shaped by taxonomic bias towards primates and, to a lesser extent, birds. This bias likely reflects both the long-standing interest in primate models for understanding the origins of human cognition and practical advantages accumulated over decades of research, including established experimental paradigms. Big-team science offers an opportunity to move beyond these historical and logistical constraints by extending standardized comparative approaches to a broader range of taxa. Teleost fishes are particularly important in this context because they offer a critical test of whether patterns of cognitive evolution established predominantly in endothermic vertebrates generalize to ectotherms, providing an important comparative perspective for understanding the processes shaping cognitive diversity across the vertebrate lineage.

Descended from a lineage that diverged from that leading to tetrapods roughly 450 million years ago, teleost fishes constitute the largest monophyletic group of vertebrates [17]. With more than 30,000 identified species to date [18, 19]—accounting for approximately 98% of all living fishes [20]—they have colonized virtually all aquatic ecosystems, an ecological diversification accompanied by a remarkable range of life-history strategies, sensory specializations, and behavioral adaptations. Such diversity provides a powerful opportunity to investigate how different traits might lead to the evolution of different cognitive systems, highlighting the potential of that group for large-scale comparative analyses. Furthermore, research on fish cognition and behavior has recently gained momentum, with the discovery that teleost fishes exhibit a range of cognitive abilities previously thought to be restricted to “higher” vertebrates like primates and other mammals—including complex forms of learning, problem solving, social cognition, and executive functions (e.g., [21–32]). Consequently, conclusions drawn primarily from primates and birds may capture only part of the processes shaping cognitive evolution across vertebrates. With their deep evolutionary divergence, extraordinary ecological diversity, and broad cognitive repertoire, fishes provide a critical and largely understudied clade to understand evolutionary processes shaping cognition across vertebrates.

Here, we extend the big-team comparative approach to fishes to investigate the evolutionary processes shaping cognitive diversity across vertebrates. The current research is part of the “ManyFishes” initiative: a large-scale collaboration of international researchers that pools expertise and resources to address key questions in fish comparative cognition and behavior [33]. In this first empirical study, called “ManyFishes 1”, we explored inhibitory control, a core executive function that encompasses different forms of behavioral regulation— from the inhibition of prepotent motor responses to more complex forms of self-control [34]— and that has been extensively studied across mammals and birds (e.g., [15, 35–38]), and more recently in reptiles [39–41], amphibians [42] and fishes (see [28, 43–45]). Although the question of whether inhibitory control reflects homologous or analogous mechanisms across animal taxa remains open, multiple lines of evidence indicate that teleost fishes are capable of inhibiting prepotent responses, for example, by suppressing immediately motivated behaviors when they are not appropriate (e.g., [28–29, 46–49]).

To assess inhibitory control, we adapted the cylinder task employed by MacLean and colleagues in their large-scale comparative study of mammals and birds [15] for use in fishes, while retaining its core design to maximize comparability. In the task, subjects must “detour” around a clear cylinder to access a food reward through its open ends, rather than approaching the visible reward directly and eventually bumping into the cylinder. Building on this comparative framework, we brought together researchers from across the globe in a coordinated effort to apply a standardized protocol across a broad diversity of teleost fishes. By pooling data collected under this common goal, we assembled the largest comparative dataset on inhibitory control in teleosts to date, spanning 22 species across 19 genera. This unprecedented scale and resolution allowed us to revisit long-standing questions in ecology and evolution by examining how phylogeny, species differences, ecology, social environment, individual characteristics, and experience contribute to variation in cognitive performance within a single standardized dataset. More broadly, this study provides a first proof of concept for a big-team approach to fish cognition, showing how global collaboration and coordinated data collection can enable comparative tests at a scale previously largely restricted to other vertebrate groups.

## METHODS

### Subjects

We tested 444 subjects representing 22 species of teleost fishes. However, due to a methodological artifact, the nine *Rhinecanthus aculeatus* individuals tested at Oxford University were excluded from the analyses (see [2]). Subjects were either individually housed or grouped with one or more conspecifics in the same aquarium throughout the experiment, as appropriate for each species. The dimensions of the aquariums differed across species (volume range: 9–320 L), with each containing 1) a resting space for fishes to hide and/or sleep (e.g., shelter) and 2) an enrichment item (e.g., ornament, plant, visual/physical access to conspecifics). Food rewards for the testing was based on the species-specific needs. The food either directly adhered to smooth surfaces (e.g., glass, acrylic, PVC) or it was blended with a sticky substrate (e.g., prawn/shrimp, gelatine, agar powder). For detailed information on the species tested and the rewards used, see **Table 1**.

**Table 1.** Information of species tested in the cylinder task. The table includes species’ scientific and common names, ecosystem (freshwater or marine), origin/history (captive-bred or wild-caught), sample size, sex (M = male, F = female, or U = unknown), age category (juvenile, subadult, adult, or unknown), housing condition during the experiment (i.e., Steps 1-4; individual or group with number of conspecifics), group size in the wild (small or large), water volume of the testing aquarium (in liters), cylinder dimensions (length x height in centimeters), food type, trial schedule, and testing location.

| Scientific name | Common name | Ecosystem | Origin | Sample size | Sex | Age category | Housing | Group size | Aquarium volume (L) | Cylinder (cm) | Food | Trials | Research group |
| --- | --- | --- | --- | --- | --- | --- | --- | --- | --- | --- | --- | --- | --- |
| *Carassius auratus* | goldfish | freshwater | captive-bred | 12 | 4M, 8F | unknown | group (3) | large | 84 | 20 x 10.16 | pellets | 10 over 3 days | Ben Gurion University (Israel) |
| *Corydoras aeneus* | bronze corydoras | freshwater | captive-bred | 15 | 7M, 8F | adult | group (10) | large | 72 | 15 x 10 | pellets | 10 over 2-3 days | University of Padova (Italy) |
| *Danio rerio* | zebrafish | freshwater | captive-bred | 17 | 10M, 7F | adult | individual | large | 16 | 5 x 4 | flakes | 10 over 5 days | University of Ferrara (Italy) |
| *Dicentrarchus labrax* | European seabass | marine | captive-bred | 21 | U | juvenile | group (10) | large (juvenile) | 120 | 11 x 4 | pellets | 10 in 1-2 day(s) | LIMIA-IRFAP, IMEDEA-CSIC (Spain) |
| *Etheostoma caeruleum* | rainbow darter | freshwater | wild-caught | 7 | 5M, 2F | adult | individual | small (solitary) | 90 | 20 x 9.8 | bloodworms | 10 over 2-3 days | University of Maryland, Baltimore County (USA) |
| *Gambusia affinis* | western mosquitofish | freshwater | wild-caught | 10 | 1M, 9F | adult | group (3) | large | 76 | 7 x 6 | flakes | 10 over 2 days | Texas State University (USA) |
| *Gambusia holbrooki* | eastern mosquitofish | freshwater | wild-caught | 22 | 12M, 10F | unknown | individual | large | 120 | 6.2 x 4.7 | flakes | 10 over 3 days | Macquarie University (Australia) |
| *Genicanthus lamarck* | blackstriped angelfish | marine | wild-caught | 7 | U | subadult | individual | small | 234 | 15 x 10 | pellets | 10 over 2 days | SEA LIFE Aquarium Kansas City (USA) |
| *Gnathonemus petersii* | peter's elephantnose fish | freshwater | wild-caught | 13 | U | adult | individual | large | 75-100 | 14 x 5.5 | bloodworms | 10 over 2-3 days | University of Bielefeld (Germany) |
| *Labroides dimidiatus* | bluestreak cleaner wrasse | marine | wild-caught | 39 | 19M, 20F | adult | group (2) | small | 60 | 10 x 6 | prawn | 20 over 2 days | Lizard Island (Australia) |
| *Micropterus salmoides* | largemouth bass | freshwater | captive-bred | 16 | U | subadult | individual | large (subadult) | 52 | 10 x 6 | bloodworms | 10 over 5 days | University of Ferrara (Italy) |
| *Neolamprologus pulcher* | daffodil cichlid | freshwater | captive-bred | 42 | 21M, 21F | adult | individual | large | 50 | 10 x 8 | flakes | 12 over 2 days | University of Bern (Switzerland) |
| *Pimephales promelas* | fathead minnow | freshwater | captive-bred | 9 | 6M, 3F | adult | group (2) | large | 14 | 8 x 4.5 | bloodworms | 10 over 2-4 days | University of Detroit Mercy (USA) |
| *Poecilia reticulata*<br>(Sample 1) | guppy | freshwater | captive-bred | 23 | 11M, 12F | adult | individual | large | 16 | 5 x 4 | flakes | 10 over 5 days | University of Ferrara (Italy) |
| *Poecilia reticulata*<br>(Sample 2) | guppy | freshwater | captive-bred | 19 | 10M, 8F, 1U | adult | individual | large | 120 | 6.2 x 4.7 | flakes | 10 over 3 days | Macquarie University (Australia) |
| *Poecilia reticulata*<br>(Sample 3) | guppy | freshwater | captive-bred | 71 | 32M, 39F | adult | group (6) | large | 9 | 5 x 4 | flakes | 10 over 3 days | Stockholm University (Sweden) |
| *Pomacentrus amboinensis* | ambon damsel | marine | wild-caught | 14 | M | adult | individual | small | 20 | 10 x 6 | prawn and flakes | 10 in one day | Lizard Island (Australia) |
| *Pseudopoecilia fria* | fria toothcarp | freshwater | wild-caught | 10 | 3M, 7F | adult | group (10+) | large | 40 | 5 x 4 | pellets | 10 over 4-5 days | Universidad de las Américas (Quito, Ecuador) |
| *Rhinecanthus aculeatus* | white-banded triggerfish | marine | wild-caught | 18 | U | adult | individual | small | 82-100 | 15 x 15 | prawn | 10 over 2 days | University of Oxford (UK), Lizard Island Research Station (Australia) |
| *Scarus iseri* | striped parrotfish | marine | wild-caught | 1 | U | subadult | individual | small | 234 | 12 x 8 | pellets | 10 over 3 days | SEA LIFE Aquarium Kansas City (USA) |
| *Scarus taeniopterus* | princess parrotfish | marine | wild-caught | 6 | U | subadult | individual | small | 234 | 12 x 8 | pellets | 10 over 3 days | SEA LIFE Aquarium Kansas City (USA) |
| *Sparisoma radians* | bucktooth parrotfish | marine | wild-caught | 2 | U | subadult | individual | small | 69–234 | 12 x 8 | pellets | 10 over 3 days | SEA LIFE Aquarium Kansas City (USA) |
| *Sparus aurata* | gilthead seabream | marine | captive-bred | 39 | M | juvenile | group (5) | large (juvenile) | 320 | 22 x 10 | pellets | 10 over 2 days | CCMAR (Portugal) |
| *Toxotes chatareus* | archerfish | freshwater (brackish) | wild-caught | 11 | U | unknown | individual | small | 84 | 20 x 10.16 | pellets | 10 over 3 days | Ben Gurion University (Israel) |

### Apparatus

Each aquarium included two “guillotine” doors, one clear and one opaque, which divided the aquarium into a holding compartment and a testing compartment (for a similar approach, see [29, 50–56]). Pulling up the opaque door followed by the clear door allowed subjects to see the set-up before having access to the testing compartment. The doors were present during parts of the acclimation phase (Steps 1-3, see below) and during the testing phase (Step 4). In all other cases, subjects were allowed to swim freely between compartments without doors.

The cylinder was made of a clear material (e.g., acrylic, PVC, glass) and was open on both sides (see **Figure 1**). The size of the cylinder was scaled proportional to the size of the species and *at least* one body length long and one body height high (estimates were based on the size of the largest individual of the species in the sample) so that subjects could enter/exit the cylinder with ease. The thickness of the cylinder was approximately 0.5 centimeters and the distance between the edge of the cylinder and the side of the tank was *at least* one body length long. The cylinder was placed in the testing compartment of the aquarium, so that the swimming distance between the doors and the cylinder was five to 10 times the body length of the fish (which varied based on the size of the aquarium).

**Figure 1.**
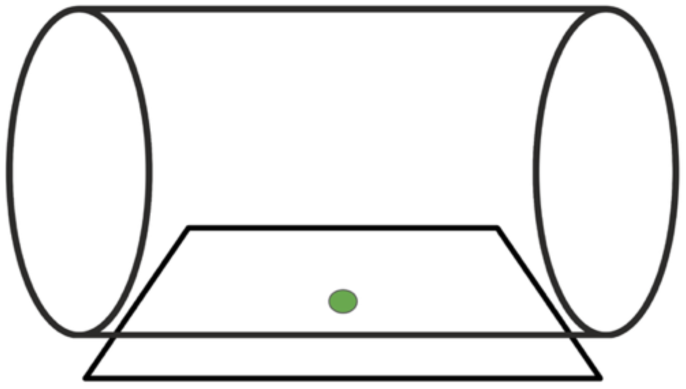
Cylinder task design. The cylinder used for testing is attached to a white base plate and contains a food reward located on a green dot drawn in the middle.

### General procedure

The protocol was based on the study by Triki and colleagues [55]. Although the design resembled largely the one used by MacLean et al. [15], it did not include the training phase with an opaque cylinder. Our experiment consisted of an *acclimation phase* (Steps 1-3), in which subjects were exposed to each component of the experiment separately prior to testing, followed by a *testing phase* (Step 4), during which data were collected. Note that housing and acclimation/testing could occur in the same aquarium or a different aquarium (e.g., for subjects living with conspecifics that needed to be separated for testing). For the testing phase, however, all subjects were prevented from visual access to conspecifics. For more details and experimenter FAQs, see **S1 Table** (Supporting Information).

#### Step 1: Plate training (doors present)

The goal of this step was to train subjects to learn to eat from a plate that mimicked part of the apparatus used for testing (Step 4). The subjects started in the holding compartment. After opening the opaque door, followed by the clear door, the fish entered the testing compartment to access a white plate presented either vertically or horizontally on the aquarium floor using weights (See **Panel A, Figure 2**). The plate was marked with a green dot located in the middle and a food reward was placed on top of the dot. Once the doors opened, they were either fully removed from the aquarium or they were kept propped open so that the fish could swim back and forth between both holding and testing compartments until they ate the food. Subjects were given a maximum of five minutes to eat the food. If they did not eat within the allotted time, the plate was removed from the compartment and a new trial began. Subjects completed Step 1 and were moved to Step 2 once they ate the food from the green dot in at least five out of six consecutive trials (i.e., > 80% success; e.g., [55, 57]). For group settings, a plate without food was made available in the common aquarium prior to starting Step 1.

**Figure 2.**
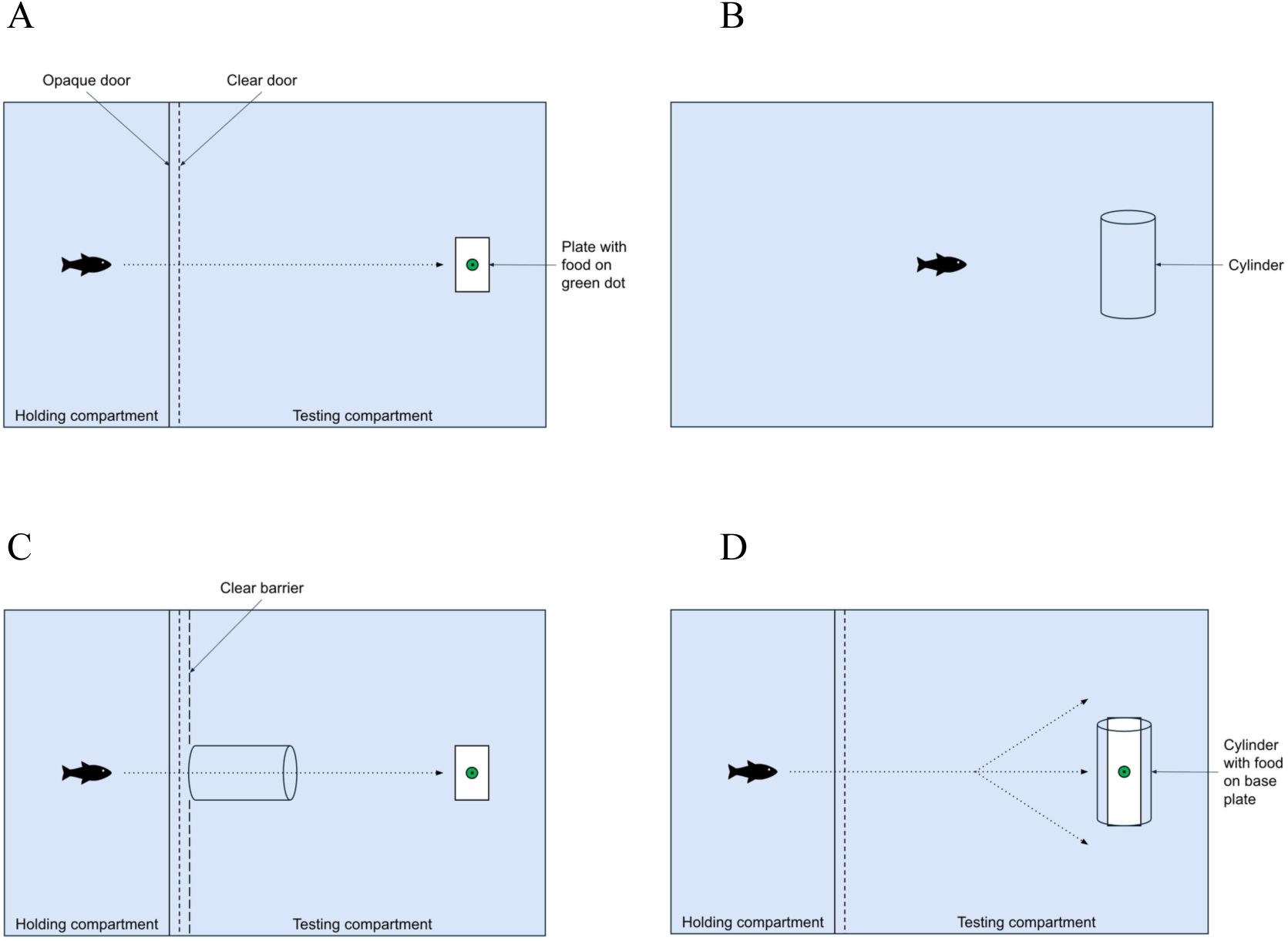
Experimental steps of the cylinder task. In Step 1 (**Panel A**), the subject starts in the holding compartment; after opening the opaque door, followed by the clear door, the subject enters the testing compartment to access a food reward located on a green dot drawn in the middle of a white plate. In Step 2 (**Panel B**), the subject is allowed to freely explore a clear cylinder for a period of 48 hours. In Step 3 (**Panel C**), the subject starts in the holding compartment; a cylinder is placed perpendicularly to the doors with clear barriers on each side so that it creates a central tunnel that the subject must pass through to access a food reward located on a green dot drawn in the middle of a white plate. In Step 4 (**Panel D**), the subject starts in the holding compartment; after opening the doors, the subject enters the testing compartment containing a cylinder attached to a white base plate that the subject must detour to access a food reward located inside it.

#### Step 2: Cylinder familiarization (doors absent)

Step 2 aimed to familiarize subjects with a clear cylinder. To achieve this, the cylinder was placed horizontally and parallel to the rear wall of the aquarium, at approximately the same distance from either side and with *at least* one body length long from each side so that the fish could swim around the cylinder with ease (See **Panel B, Figure 2**). Subjects were allowed to freely explore the cylinder in the testing compartment for a duration of 48 hours. For group settings, a single cylinder was placed in the common aquarium.

#### Step 3: Forced cylinder trial (doors present)

Prior to testing, subjects were required to enter and exit the cylinder to familiarize them with the apparatus and facilitate interaction with it. Subjects started in the holding compartment. A white plate carrying a food reward was placed in the testing compartment and a cylinder was placed perpendicularly to the doors—which were blocked by clear barriers—to create a central tunnel that subjects had to enter and exit to reach the food (See **Panel C, Figure 2**). Additional food rewards could be placed within the testing environment to encourage subjects to enter and exit the cylinder. If the subject did not enter the cylinder within five minutes, the plate and cylinder were removed from the compartment and a new trial began. Subjects completed Step 3 and moved to Step 4 if they entered and exited the cylinder *once* (whether they ate the food or not).

#### Step 4: Testing (doors present)

Step 4 aimed to test subjects in the cylinder task. Subjects started in the holding compartment. Before the testing began, each subject received one plate trial (equivalent to a “warm-up” trial). Testing started with the experimenter placing the cylinder containing a food item in the middle— now attached to a white base plate (See **Panel D, Figure 2**)—in the testing compartment. The plate base aimed to guarantee stability underwater and consisted of a white flat rectangle (or half tube) with metal weights. The amount of food placed in the cylinder was equivalent to approximately the volume of food which would fill the subject’s mouth when foraging (one “bite”). A trial began when the experimenter pulled up the opaque door followed by the clear door (approximately two seconds total), thereby allowing the subject to enter the testing compartment and interact with the cylinder. To reach the food reward, the fish had to “detour” the cylinder walls and swim inside the cylinder to retrieve the food located on the green dot. The doors were removed and/or kept propped open during the entire trial. Subjects received at least 10 trials over two to four consecutive days, with a minimum of three trials per day. Trials were conducted with a minimum intertrial interval of 10 minutes, but more typically the duration of testing all subjects for one trial.

For each trial, the experimenter reported whether the subject 1) ate the food without touching the cylinder (noted as “*Success*”) or 2) touched the cylinder before eating *or* without eating the food (noted as “*Failure*”), within five minutes. Trials in which the subject did not touch the cylinder and did not eat the food within the allotted time were still reported (noted null response or “NA”) but they were not used for the main analysis. Performance in the task was measured as the proportion of successes vs. failures.

### Data analysis

Analysis was run in RStudio (version 2022.07.2 Build 576; Integrated Development for R. RStudio, PBC, Boston, MA, U.S.A.). The statistical threshold was set at P = 0.05.

#### Question 1. Does phylogenetic relatedness explain cognitive performance?

We tested whether species with closer phylogenetic relationships performed more similarly in the task than more distantly related species [13, 59]. We estimated this resemblance using the phylogenetic signal measured in Pagel’s lambda (λ)—a measure of similarities in traits across species given a shared evolutionary history— here quantifying the strength of the association between similarity in task performance and phylogenetic relatedness [60]. Values of λ typically range from 0 to 1, with lower values indicating that trait variance is weakly associated with phylogeny. It is important to note that estimates of phylogenetic signals may be unreliable when sample sizes are small (i.e., fewer than 30 species; [61]), potentially biasing λ estimates.

However, we adopted Pagel’s lambda as simulation studies reveal that it produces the lowest rates of type I and type II errors when estimating true phylogenetic signals, compared to the most common other indices (see [60]). Statistical support for the presence of phylogenetic signals (i.e., λ value significantly greater than zero) is commonly evaluated using likelihood ratio tests.

To estimate phylogenetic signal, phylogenetic distances were extracted from Betancur-R et al. (2017)’s study [62], a widely used and up-to-date phylogenetic classification of bony fishes. When data for a focal species were unavailable, we used data from a congeneric species; this was the case for *Etheostoma caeruleum* (represented by *E. vitreum*), *Neolamprologus pulcher* (*N. brichardi*), *Pimephales promelas* (*P. vigilax*), *Poecilia reticulata* (*P. latipinna*), *Pomacentrus amboinensis* (*P. brachialis*), and *Sparisoma radians* (*S. viride*). Note that *Scarus iseri* and *Scarus taeniopterus* were combined into the genus *Scarus*, and *Gambusia affinis* and *Gambusia holbrooki* were combined into the genus *Gambusia*. Because phylogenetic distances for *Carassius auratus*, *Corydoras aeneus*, and *Genicanthus lamarck*, were unavailable, we used the closely related genera/species *Cyprinus carpio*, *Callichthys callichthys*, and *Centropyge bicolor*, respectively, for the analysis. Note that *Pseudopoecilia fria* was excluded from the analysis because its genus was not represented in the classification. For *Poecilia reticulata*, *Scarus*, and *Gambusia*, performance was calculated as the average across all subjects and research groups.

To obtain performance for each group, we aggregated the data across trials for each subject—i.e., the number of successful trials divided by the total number of trials—and then computed the average performance per genus. We tested the significance of phylogenetic signals using the “phylosig” function from the “phytools” R package [63]. We also performed Akaike Information Criterion (AIC) model selections by comparing our data with a Brownian motion model of cognitive evolution, in which cognitive variance among species is directly related to the length of shared evolutionary history (referred to as λ = 1), and a null model, with no significant relationship between cognitive variance and phylogenetic distance (referred to as λ = 0).

#### Question 2. Does species predict performance?

To estimate the effect of species on cognitive variance, we analyzed the binomial cognitive responses using a generalized linear mixed-effect model (hereafter GLMM) with binomial distribution fitted with species (category), age category (juvenile, subadult, adult), sex (female, male, unknown), ecosystem (freshwater, marine), origin/history (captive-bred, wild-caught), and group size (small, large) as fixed factors, trial number as covariate, and subject ID as random effect. We used the “glmer” function from “lme4” R package [64], specifying “optmix” as control parameter optimizer and “bobyqa” as the algorithm for the optimization control setting [65]. We assessed the relevance of species via AIC model comparison (“anova” function from “car” R package) by fitting a second model without this predictor. The significance of the model parameters was assessed via Wald chi-squared method (“Anova” function from the “car” R package; [66]). We also compared the homogeneity of variance on the aggregated performance across species using the “LeveneTest” function from “car” R package. The dataset used for this analysis consisted of the first 10 trials completed by each subject, with missing data (NAs) excluded and trial numbers adjusted accordingly. Only subjects that completed at least two trials were included in the analysis, thus excluding individuals who did not complete any trials (i.e., all NAs) or who completed one trial only.

#### Question 3. What other possible factors impact performance?

The relevance of age category, ecosystem, origin/history, sex and group size on cognitive performance was assessed using a GLMM fitted with those terms as fixed factors, and subject ID and species as random effects. The model also included trial number (continuous variable) as predictors. The model included the interaction term species × trial number because we expected between-species difference on the cognitive improvement across the 10 trials. We also included the interaction term species × age to assess age differences in the cognitive performance within species. Parameter estimation was assessed via Wald chi-squared method. The analysis was based on the same dataset used for Question 2. The initial model included the two interaction terms showed high multicollinearity between predictors (adjusted VIFs > 5.91). By inspecting the model matrix, we also observed rank-deficient due to lack of observed dummy variables depending on the contrasts mainly defined by the interaction terms. Therefore, we limited this first analysis by using a model fitted with main predictors without interaction terms. Then, we specifically estimated the significance of trial, sex, and research group, by running three different GLMMs on different restricted datasets (see below). For the analyses on the effect of sex on performance, see Supporting Information.

#### Question 4. Does performance increase due to learning?

The GLMM used for further analyzing the effect of trial number was fitted with trial number as covariate, species as fixed factor, and subject ID as random effect. The model included the interaction between trial number and species. Model selection was performed via comparing AIC between the null model (i.e., without trial as covariate) and the model without the interaction term. The analysis was based on the same dataset used for Question 2.

#### Question 5. Case study: Do guppies tested across three research groups perform similarly?

We used research institution as a proxy for variation in housing conditions for *P. reticulata*, which was tested by three independent research groups. We tested the potential research group effect on cognitive performance by comparing two GLMM models, one with and the other without this term as a fixed factor.

## RESULTS

### Descriptive statistics

We report the summary statistics for each species included in the cylinder task in **Table 2** and **Figure 3**.

**Table 2.**
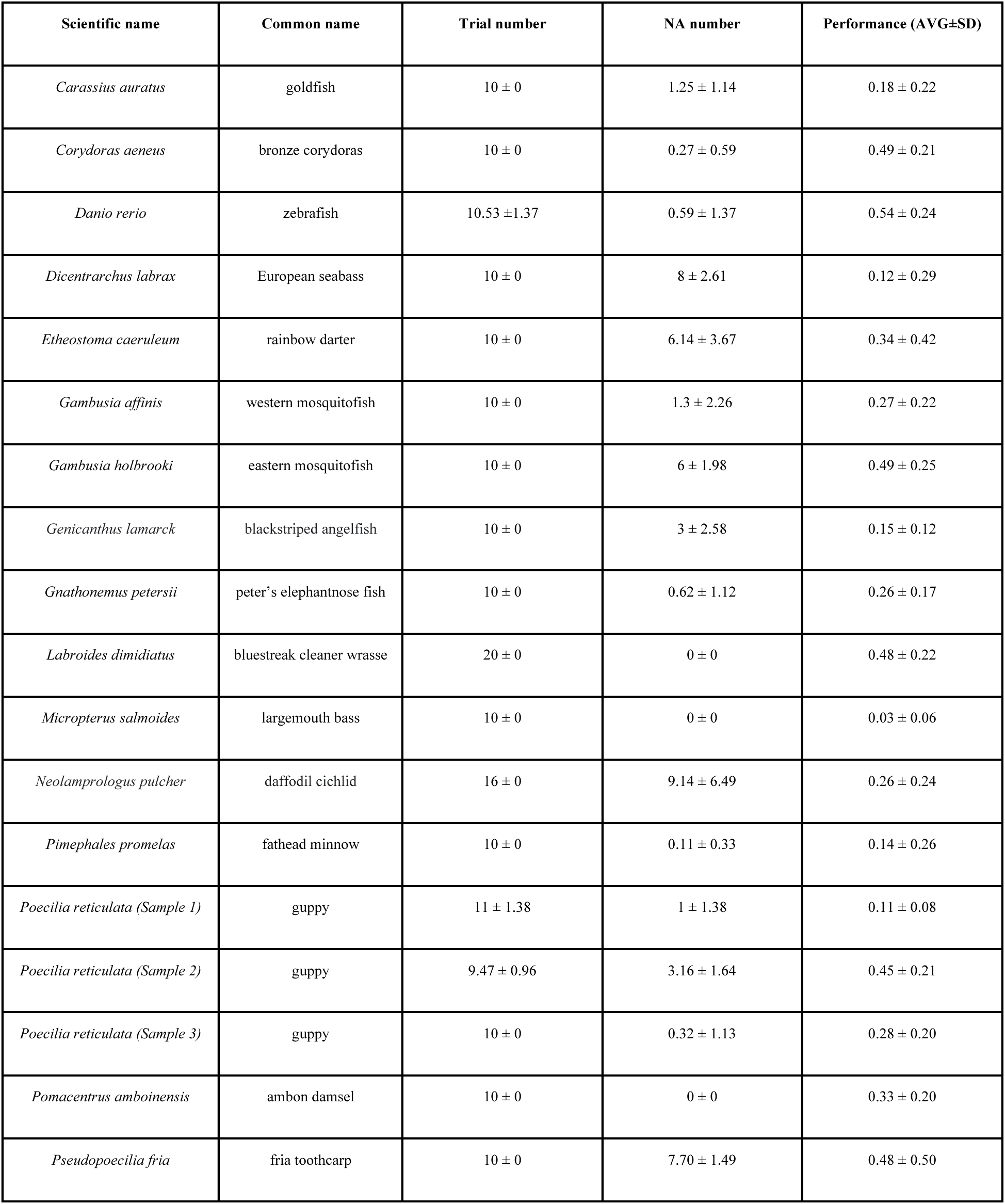
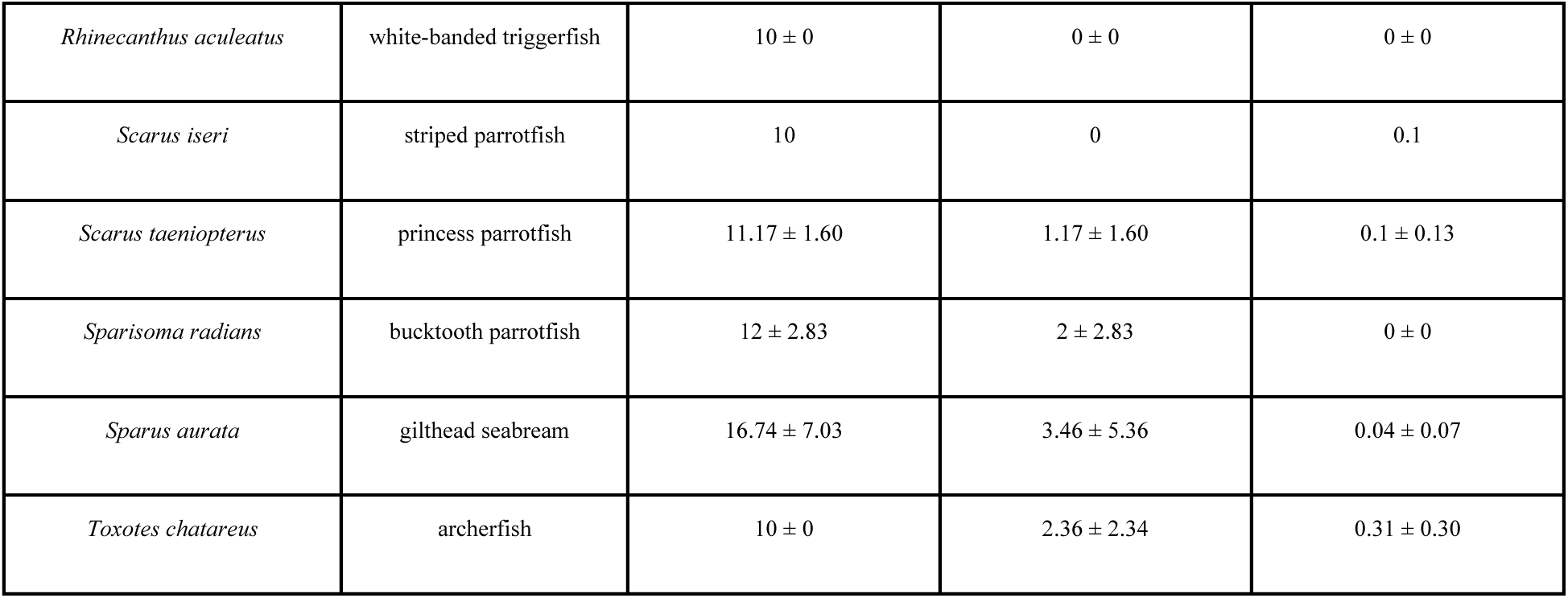
Average (± SD) species performance in the cylinder task, including the number of trials completed, number of null responses (NA), and proportion of successful trials (performance), calculated from subjects that completed at least 10 trials.

**Figure 3.**
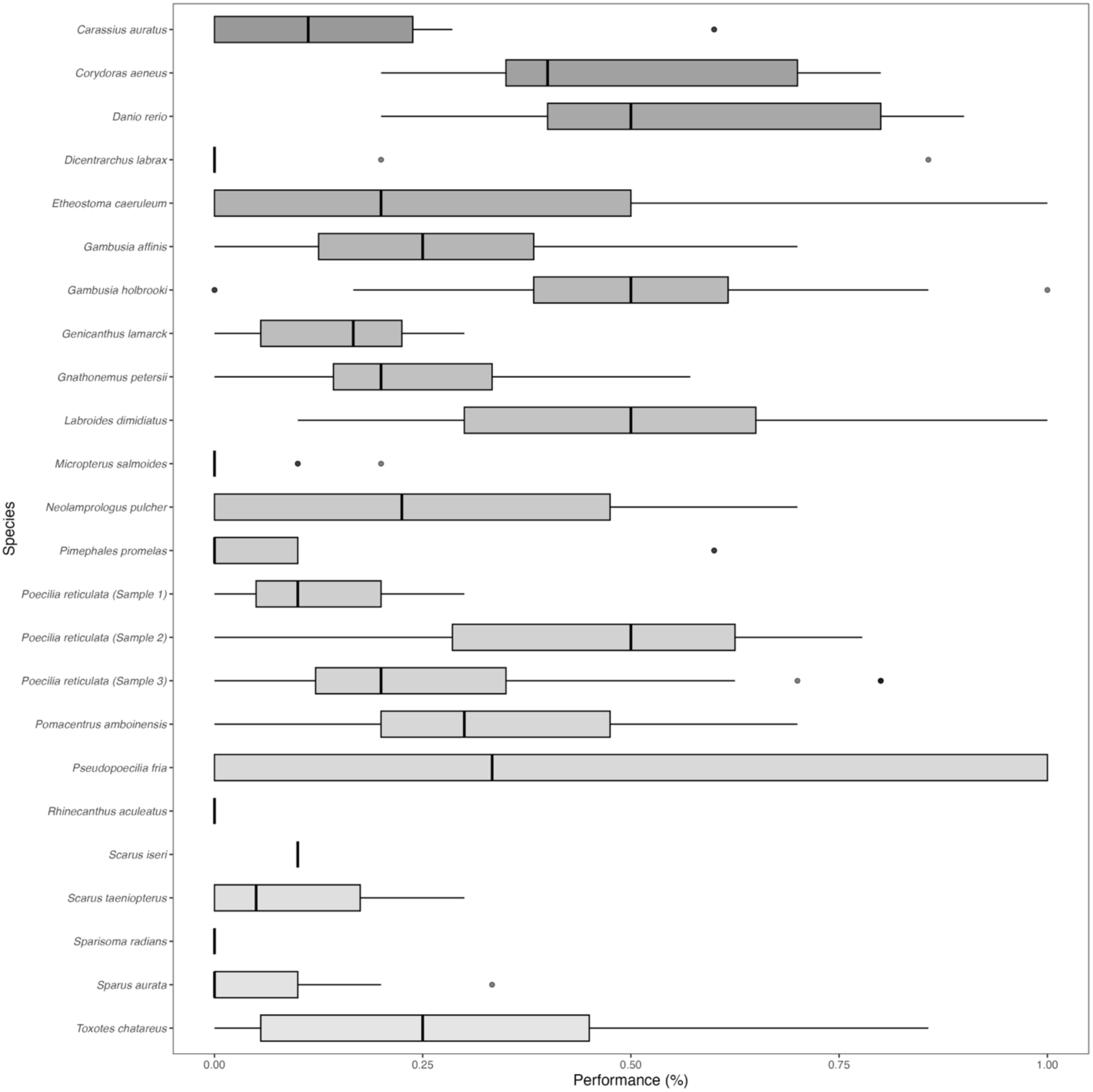
Boxplots of species’ proportion of successful trials (performance) in the cylinder task.

### Result 1. Phylogenetic relatedness did not explain performance

The phylogenetic analyses showed a value of phylogenetic signal λ that did not differ from zero (maximum likelihood estimate of Log λ = -80.40, P > 0.99; **Figure 4**). The best model in explaining the data (i.e., lower AIC value via maximum likelihood estimation [67]) was the model that did not account for the phylogenetic signal (no signal model: λ = 0, AIC: 165.543), compared to the model that included the phylogenetic data (AIC: 168.393), or the Brownian model (AIC: 171.829).

**Figure 4.**
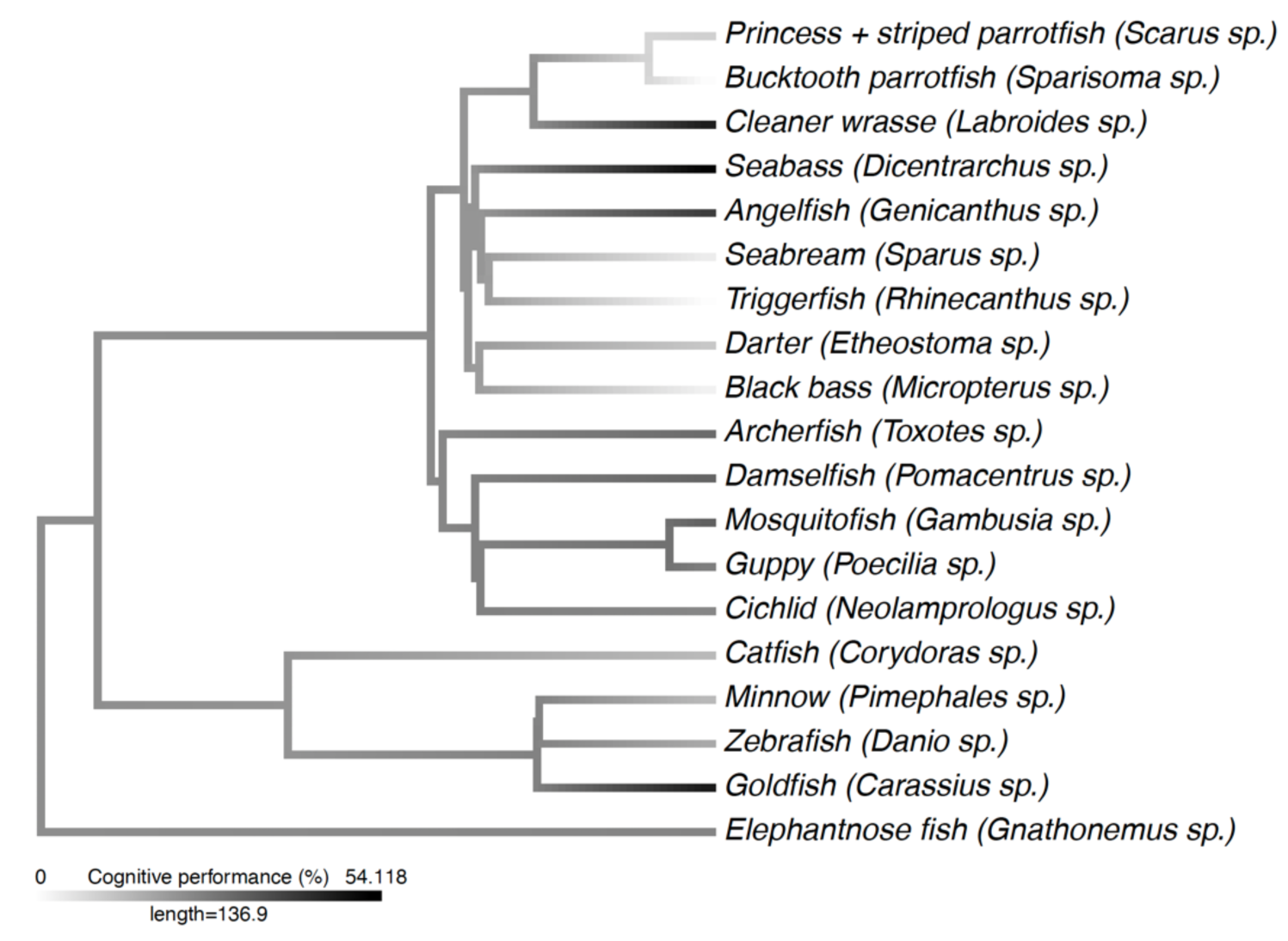
Phylogenetic analyses (signal λ) across 19 fish genera tested in ManyFishes 1. Phylogenetic data retrieved from Betancur-R et al. (2017)’s study [62].

### Result 2. Species and individuality explained variation in performance

The model fitted with species as fixed effect and origin/history, sex, age category, ecosystem, group size, and trial number as fixed effects, revealed a significant effect of species on performance (χ^2^_15_ = 73.201, P < 0.001), but the model failed to achieve convergence likely attributed to the data (i.e., different number of responses per species), consequently producing weak effect estimation. Using species as random effect—an approach that is more conservative (see [13])—and origin/history, sex, age category, ecosystem, group size, and trial as fixed effects, we found that the full model was a better fit to our data than either the null model or the model that did not include species (LRT, χ^2^_1_ = 42.119, P < 0.001). Indeed, a comparison of the AIC indicated that the full model (AIC: 3540.1) better minimized information loss than the null model (AIC: 3542.8) or the model without species (AIC: 3580.3). We evaluated species differences by extracting and plotting random effects (**Figure 5**). A test comparing homogeneity of variance across species indicated that some species showed significantly larger interindividual variance than others (Levene’s Test, F_21,380_ = 4.32, P < 0.001; **Figure 6**).

**Figure 5.**
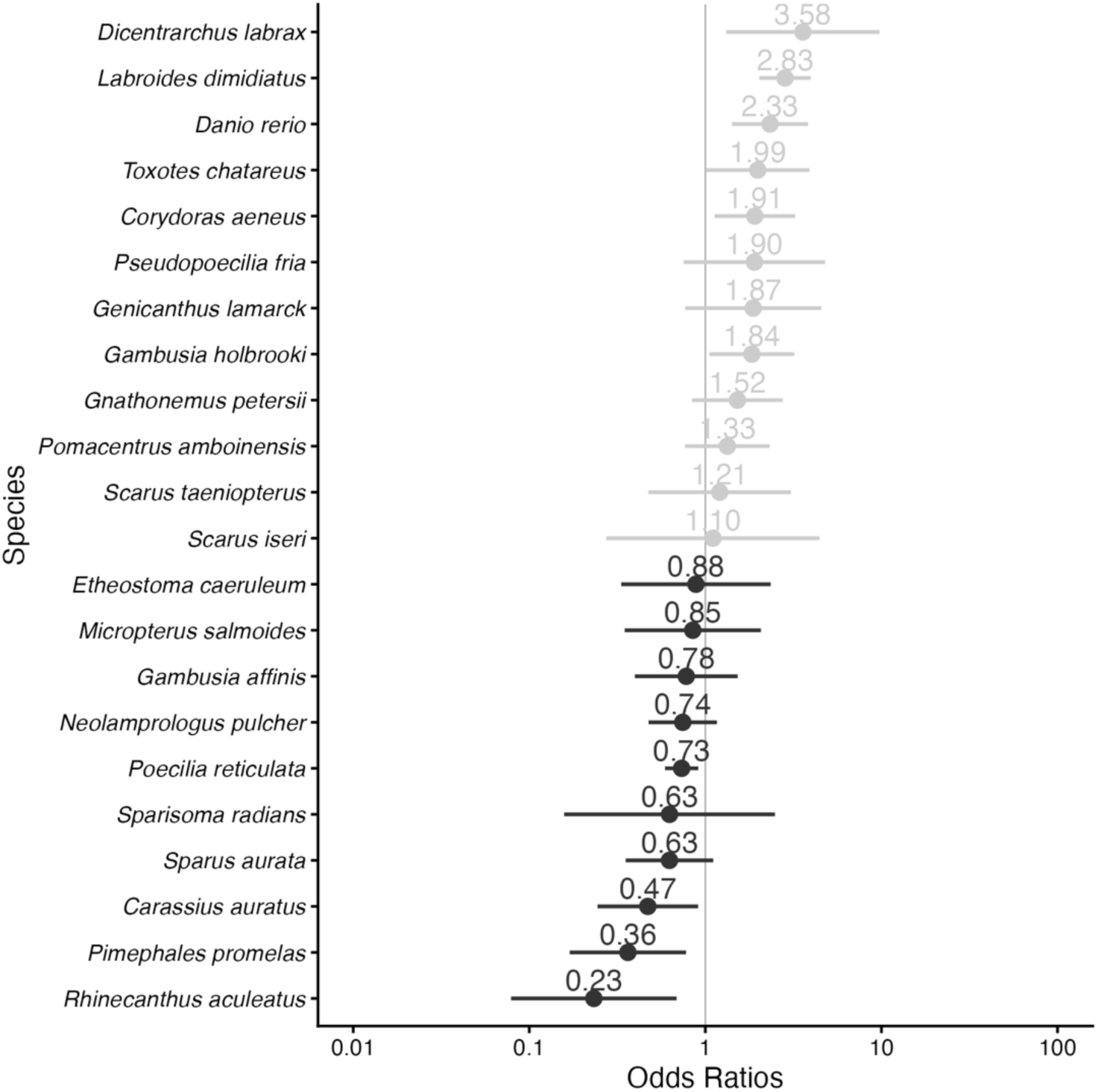
Forest plots for GLMM representing odds ratios for species random effects in the model. The figure shows how each species’ intercept differs from the average intercept. Higher odds ratios (i.e., right of the grey line) indicate better performance.

**Figure 6.**
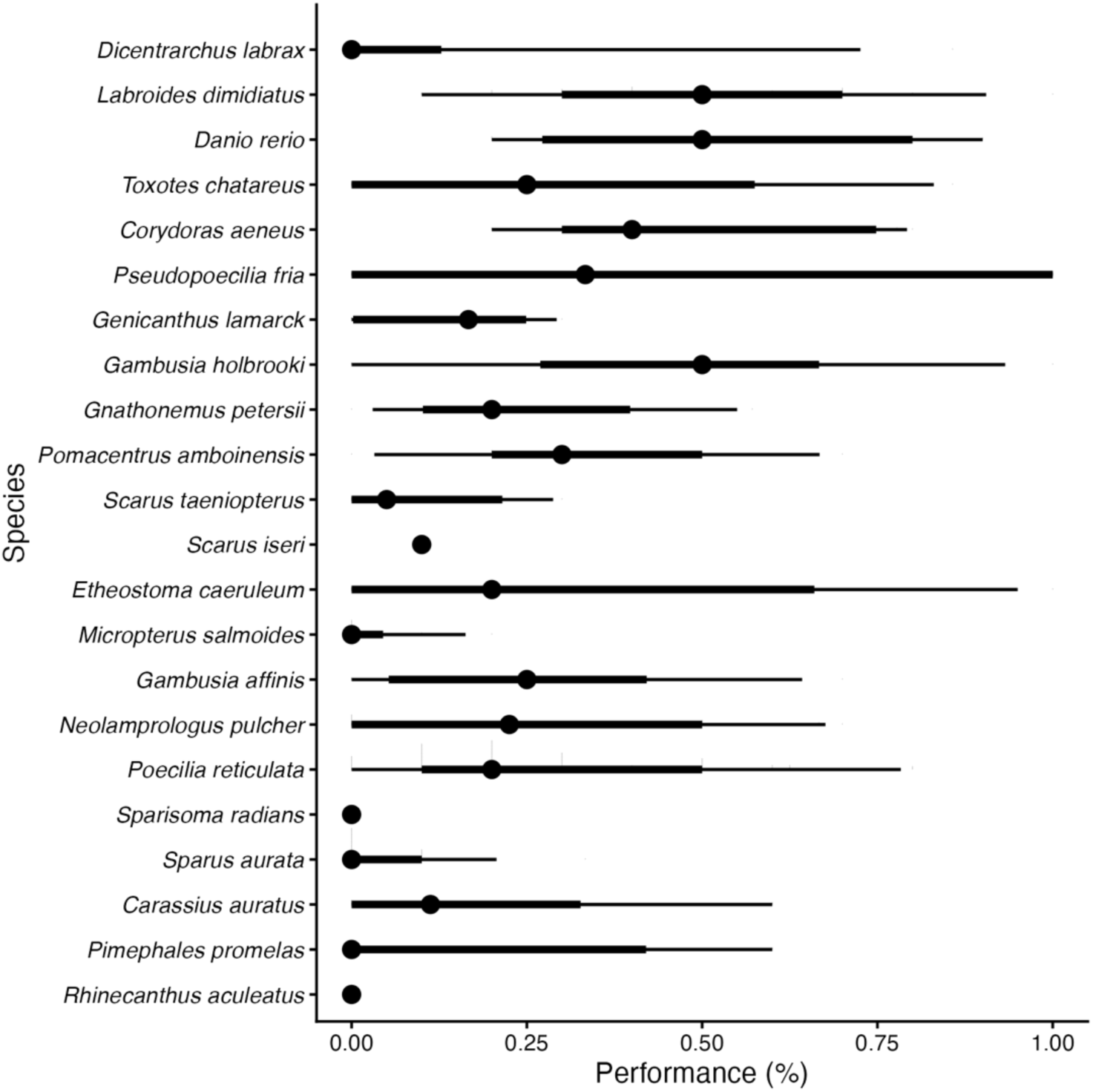
Quantile dotplot-interval representing the distribution of variance across species. The central dot per species represents the mean performance of the distribution. The thick line indicates the higher-density interval with a 66% probability of observing the data within this range, whereas the thin line indicates the lower-density interval with a 95% probability range.

### Result 3. No other tested factors influenced performance

Subjects’ performance was not explained by any of the factors included in the model (age category: χ^2^_2_ = 4.491, P = 0.11, sex: χ^2^_2_ = 2.941, P = 0.23; origin/history: χ^2^_1_ = 0.021, P = 0.88; ecosystem: χ^2^_1_ = 0.098, P = 0.75; group size: χ^2^_1_ = 0.083, P = 0.77; trial: χ^2^_1_ = 2.607, P = 0.11). A closer examination of the forest plots indicate a potential effect of ontogeny, with adults outperforming juveniles in the task (but this was not captured by the model; **Figure 7**).

**Figure 7.**
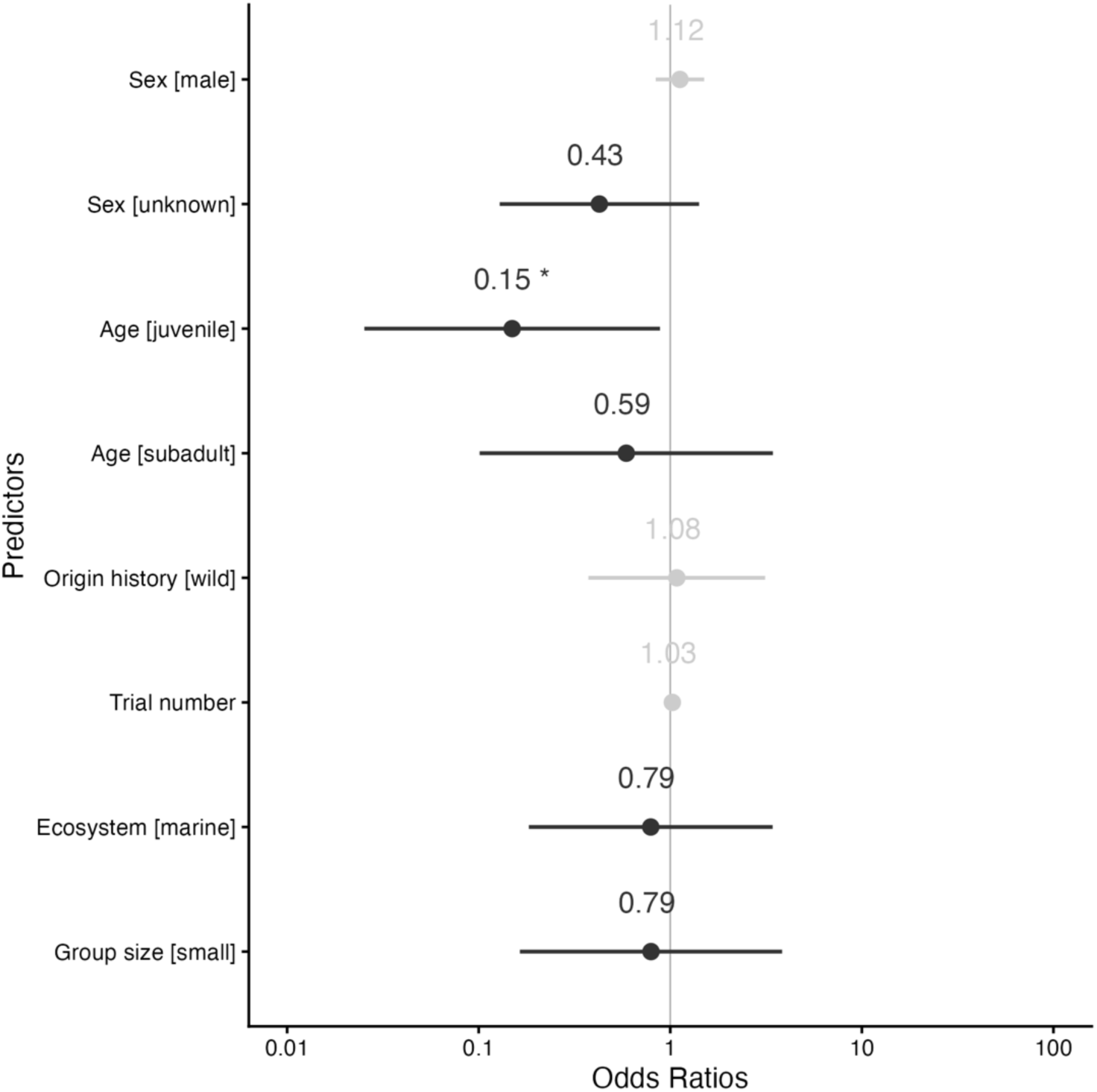
Forest plots for GLMM representing odds ratios for fixed effects in the full model that included sex, age category, origin/history, trial number, ecosystem, and group size, as fixed effects. Higher odds ratios (i.e., right of the grey line) indicate better performance.

### Result 4. Learning affected performance differently across species

The model revealed a significant interaction between species and trial number (χ^2^_21_ = 47.66, P < 0.001; **Figure 8**). The full model with the interaction (AIC: 3489.1) was a better fit to the data than either the null model (AIC: 3706.6) or the model that did not include the interaction (AIC: 3500.1), suggesting interspecific differences in learning.

**Figure 8.**
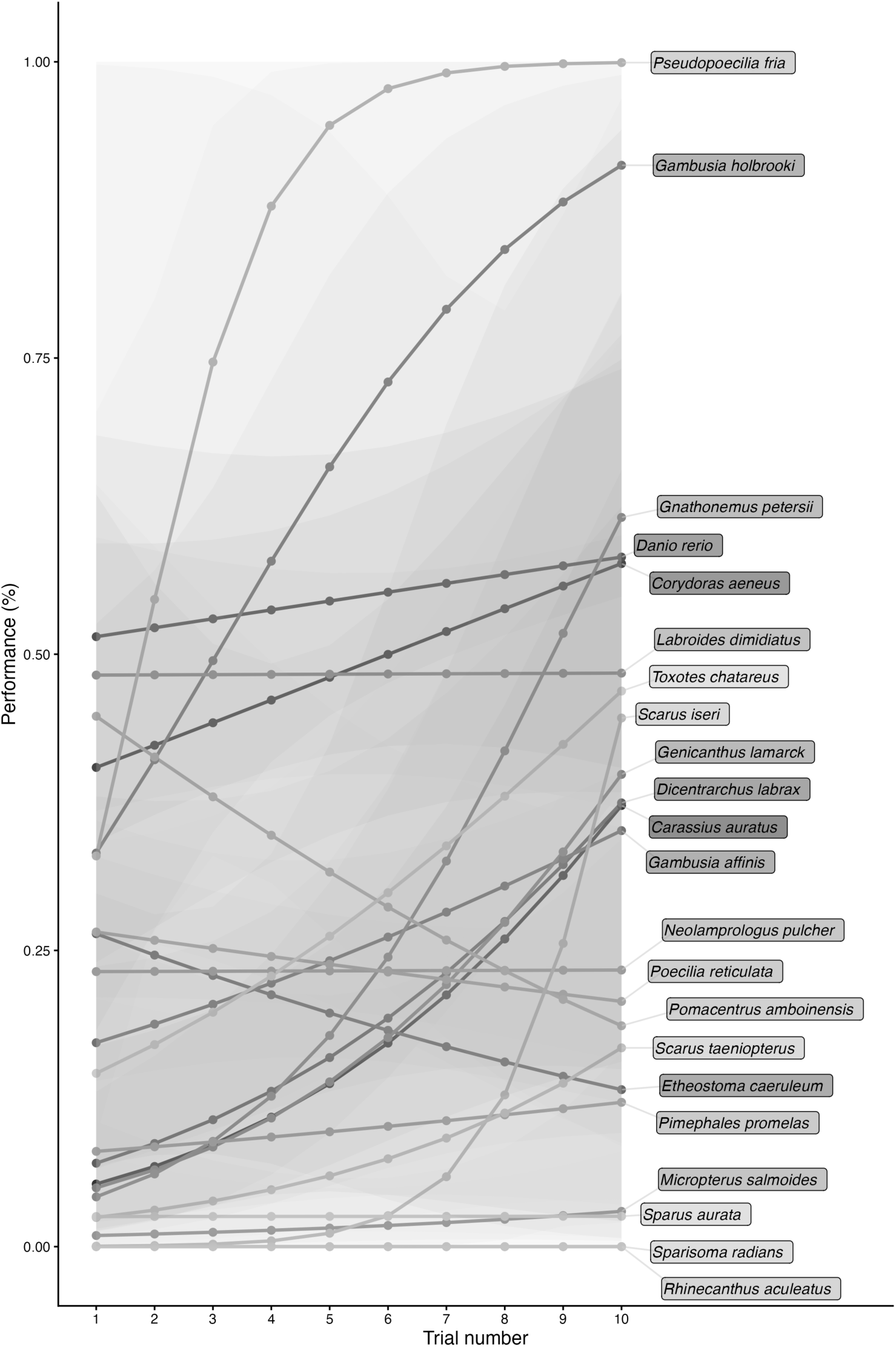
Species’ proportion of successful trials (performance estimates) across trials. Note that *Scarus iseri* and *Sparisoma radians* were excluded from the analysis.

### Result 5. Case study: Guppies tested by three research groups performed differently

When comparing the three samples of *Poecilia reticulata*, the model revealed a significant effect of research group on performance (χ^2^_2_ = 36.39, P < 0.001; **Figure 9**). Specifically, subjects at Macquarie University performed better than those at Stockholm University and University of Ferrara (MU-SU: estimate difference = 0.847, se = 0.261, P = 0.003; MU-UF: estimate difference = 2.043, se = 0.341, P < 0.001), whereas subjects at Stockholm University performed better than those at University of Ferrara (SU-UF: estimate difference = 1.196, se = 0.269, P < 0.001). The full model (AIC: 1132.0) was a better fit to the data than the null model (AIC: 1163.0).

**Figure 9.**
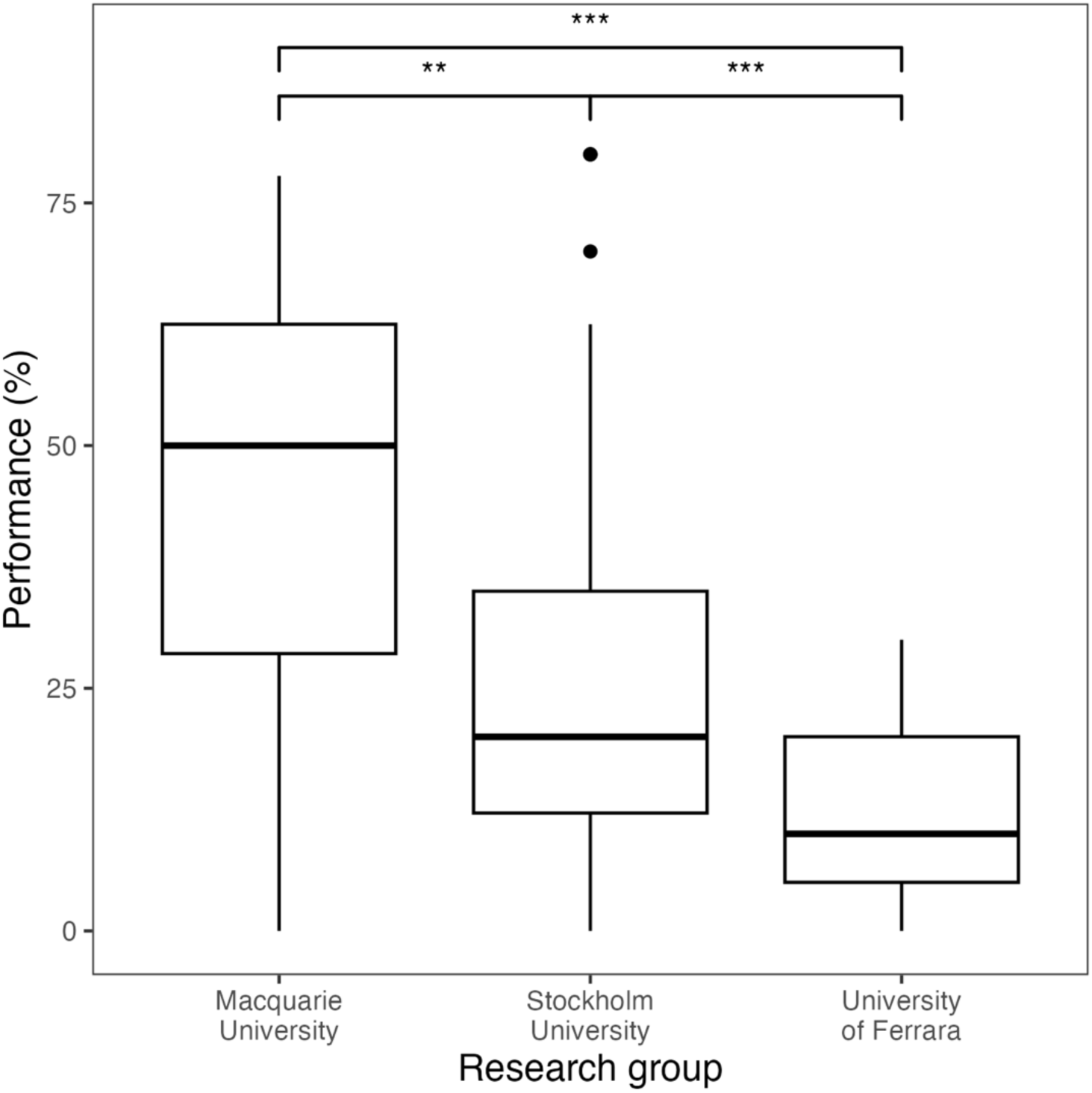
Boxplots of *Poecilia reticulata*’s average proportion of successful trials in the cylinder task performed by three different research groups.

## DISCUSSION

In this study, we asked whether big-team science could bring comparative cognition to an often overlooked vertebrate group—teleost fishes—by bringing together researchers across the globe to apply a standardized cognitive task across a variety of species. As part of the ManyFishes collaboration, we were able to gather 15 research groups from 10 countries that successfully collected inhibitory control performance data in the classic cylinder detour task from 444 individuals across 22 species of teleost fishes representing 19 genera. To our knowledge, this represents the largest single dataset on comparative cognition in fishes to date, and a sample size comparable to those of other big-team science studies. Our analyses revealed substantial variation in performance both between *and* within species, accompanied by a weak phylogenetic signal—a finding that deviates markedly from previous research in mammals and birds.

Although there was no significant overall effects of age, ecosystem, origin/history, sex, and group size, performance changed differently across species as a function of trial number, indicating important interspecific differences in learning patterns. Below, we discuss these findings and their potential implications for our understanding of vertebrate cognitive evolution.

The lack of significant phylogenetic signal and the substantial variability in task performance is interesting for several reasons. Previous studies in mammals and birds reported much higher phylogenetic signals, albeit not always significant and largely driven by datasets that are heavily skewed toward primates and comparatively younger and more tightly sampled clades than teleost fishes (e.g., [12–16]). Assuming that inhibitory control is homologous within teleost fishes, our findings may indicate that the trait has been repeatedly and markedly shaped by evolutionary processes among closely related lineages. Closely related species exposed to different ecological pressures may therefore rapidly evolve different inhibitory control performance, whereas eventual similarities may be more likely ascribable to similar selective pressures and convergent evolution. Notably, although this pattern was not observed across all species, similarly high performance in the cylinder task was often associated with species experiencing strong ecological pressures in the wild, such as small-bodied prey species (e.g., *Danio rerio*, *Gambusia holbrooki*) or specialized foragers (e.g., *Labroides dimidiatus*). Alternatively, the large interspecific variation combined with a low phylogenetic signal may also emerge in case of traits that display highly phenotypic plasticity, which has been often observed for cognitive traits in fishes [68, 69]. These aspects warrant further investigation in future studies.

The large interindividual variation in performance is another notable finding and may partly account for the weak phylogenetic signal we observed. Individual differences in fish cognition are increasingly recognized [70], and previous work on inhibitory control has reported substantially greater interindividual variation in fishes than is typically observed in other vertebrates [71]. Such variation may reflect differences in developmental history, experience, and rearing conditions [54, 72–78], but may also be compounded by unavoidable differences among testing sites and subtle variation in task implementation [79]. Indeed, aquarium structure was recently suggested to influence cylinder task performance in triggerfish tested across two sites [2]. Our repeated sampling of *P. reticulata* across three research groups provides further evidence that performance may vary across testing environments, despite the use of a standardized protocol. Thus, the substantial interindividual variation observed here may reflect a combination of inherent biological differences and heterogeneity associated with testing environments and methodological implementation.

More broadly, the combination of substantial interindividual and interspecific variation feed to the divide between endotherm and ectotherm vertebrates. Endotherms such as birds, for example, possess relatively larger brains than ectotherms but exhibit comparatively limited intraspecific variation, potentially reflecting by the energetic support provided by parental care (see [80]). Teleost fishes, in contrast, exhibit greater neural plasticity, in part due to their capability for continuous neurogenesis throughout life [81]. This capacity may allow fishes—and potentially other ectotherms such as reptiles and amphibians—to adapt to the ecological challenges they face (both environmental and social) by selectively growing relevant neurons in brain regions with distinctive functions. Although endotherms may exhibit similar forms of plasticity, these processes may operate at different rates or under different constraints. Consistent with this claim, brain allometry in the form of brain-body regression slopes in adults are steeper in ectotherms than in endotherms—i.e., as fishes grow in body size, their brains increase in size as well, unlike in endotherms [82–83] (see [84]). As a result, differences in brain organization at morphological, developmental, and cellular levels, together with high levels of phenotypic plasticity, may contribute to greater variation in cognitive performance in teleost fishes—both within and among species—than is typically observed in mammals and birds. Such differences raise the possibility that cognitive systems in ectotherms may exhibit larger evolvability overall than those of endotherms.

Another noticeable finding was the interaction between species and trial number. Although similar effects have been previously reported across vertebrates (e.g., [29, 85, 86]), in our study, changes in performance over trials varied among species, with some showing steep improvement, others showing only slight apparent improvement, and still others showing little or no improvement. This pattern suggests that different species may rely on different decision rules when facing the task. In some species, learning processes associated with repeated trials may have been co-opted more strongly to solve the inhibitory control task. Future ManyFishes studies, and big-team science more broadly, could further investigate species differences in cognitive performance by generating species-specific predictions about the potential role of learning.

Finally, our findings should be viewed in light of potential methodological considerations that may have influenced the inferences that can be drawn from our results. First and foremost, although this first ManyFishes study provides the largest comparative dataset of cognitive performance in fishes to date, its opportunistic nature resulted in uneven and limited phylogenetic coverage (especially relative to the number of identified fish species to date), which may have reduced our ability to detect phylogenetic patterns. This illustrates both the challenges and strengths of big-team comparative cognition: standardized multi-site approaches greatly reduce the methodological heterogeneity inherent in traditional cross-study comparisons, but cannot eliminate population- and site-specific effects. Expanding both taxonomic coverage and the replication of species across research facilities should help disentangle these sources of variation [10]. Big-team science should therefore be viewed not as a replacement for controlled laboratory studies, but as a complementary approach for identifying broad comparative patterns that can subsequently be examined mechanistically at finer biological scales.

Additionally, the finding that overall performance in fishes was lower than that recorded in MacLean et al.’s study for mammals and birds [15] could be explained by the fact that we opted for a more stringent version of the procedure. Indeed, we eliminated the initial training phase with an opaque cylinder before transitioning to a clear cylinder, as was done previously. The rationale for this change was to minimize the potential influence of prior learning of the detour route [87] and the formation of habits to the reward [88], thereby allowing us to assess inhibitory control of fishes’ impulses more directly with the reward being visible. Supporting this argument, Lucon-Xiccato and colleagues [29] used a design similar to that of MacLean et al.’s study, with guppies (*P. reticulata*), and found that their performance was comparable to that of mammals and birds. A quick comparison of performance across studies revealed that guppies from Lucon-Xiccato et al. scored generally higher than those from the current study, despite one sample being tested by the same research group. Therefore, route planning and habit formation are other potential confounding mechanisms driving the performance recorded in a test supposedly evaluating an executive function ability.

In conclusion, ManyFishes 1 provides the first large-scale assessment of cognitive performance in teleost fishes. Our results demonstrate that a common cognitive task can be successfully implemented across taxonomically and physiologically diverse species, while also revealing substantial differences among species in the extent of performance variation. This variability likely reflects a combination of evolutionary divergence, phenotypic plasticity, and methodological influences. By providing a broad comparative framework, our study creates opportunities for fine-scale investigations of cognitive variation among closely related species with contrasting ecological traits. More broadly, our findings suggest that understanding cognitive evolution across vertebrate lineages requires disentangling the contributions of genetic differences among species from the processes that shape the expression of cognitive phenotypes during ontogeny. The relative importance of these two sources of variation may differ fundamentally between endotherms and ectotherms, potentially providing a key to understanding why broad comparative patterns of cognition in fishes differ from those observed in mammals and birds.

## ACKNOWLEDGMENTS

We thank Kayla Leyden, Hannah Miller, and all the staff at SEA LIFE Aquarium Kansas City for helping with the experiments on angelfish and parrotfish. We also thank Niclas Kolm and Elli Argyriou for their contributions to data collection on guppies, Angelo Guadagno and Barbara Taborsky for African cichlids, and Marine Amann and Corentin Lebet for cleaner fish and damselfish. We are also grateful for the logistical and technical support provided by Stockholm University, the Ethological Station at the University of Bern, and the directors and staff of Lizard Island Research Station. Artificial intelligence (AI) tools were used solely for language improvement and readability, without generating original content.

## AUTHOR CONTRIBUTIONS

The authors of this work are listed alphabetically with the exception of Laurent Prétôt (first author), who coordinated the study and manuscript preparation as lead of the ManyFishes 1 collaboration; Elia Gatto (second author) and Tyrone Lucon-Xiccato (third author), who made key contributions to the data analyses and writing; and Zegni Triki (last author), who provided extensive review and editing of the final version. Authorship was determined following the ManyFishes Collaboration Agreement guidelines, which follow an adapted CRediT taxonomy.

## ETHICS STATEMENT

All experiments were performed in accordance with relevant institutional and national guidelines and regulations. It was the responsibility of each primary investigator to ensure that the appropriate ethical approval was obtained prior to data collection in relation to this study.

## DATA AVAILABILITY

Upon acceptance, the full dataset and analysis code will be made publicly available via an open-access repository.

