## Supporting Information for "Big-team science reveals that patterns of inhibitory control variation in teleost fishes differ from those in mammals and birds"

### Data analysis

***Is performance contingent on sex?***

The GLMM used for assessing the relevance of sex between *and* within species was fitted with sex and species as fixed factors, the interaction term, and subject ID as random effect. Only species with subjects from both sexes were included for the analysis. The species *Dicentrarchus labrax*, *Genicanthus lamarck*, *Gnathonemus petersii*, *Micropterus salmoides*, *Rhinecanthus aculeatus*, *Scarus iseri*, *Scarus taeniopterus*, and *Sparisoma radians* were excluded from the analysis because sex information was unavailable. In addition, the species *Pomacentrus amboinensis*, *Sparus aurata*, and *Toxotes chataerus*, were excluded because subjects belonged to a single sex.

### Results

***Is performance contingent on sex?***

The model fitted with species and sex did not reveal an effect of sex ($\chi^2_1$ = 1.62, P = 0.20) nor an interaction between these two terms ($\chi^2_{10}$ = 10.21, P = 0.42; **S1 Figure**). Although the models' comparison did show a significant difference between the model with and without the interaction term ($\chi^2_{10}$ = 19.847, P = 0.03), the boxplots seem to indicate significant sex differences for some species (but those were not picked up by the model; **S2 Figure**).

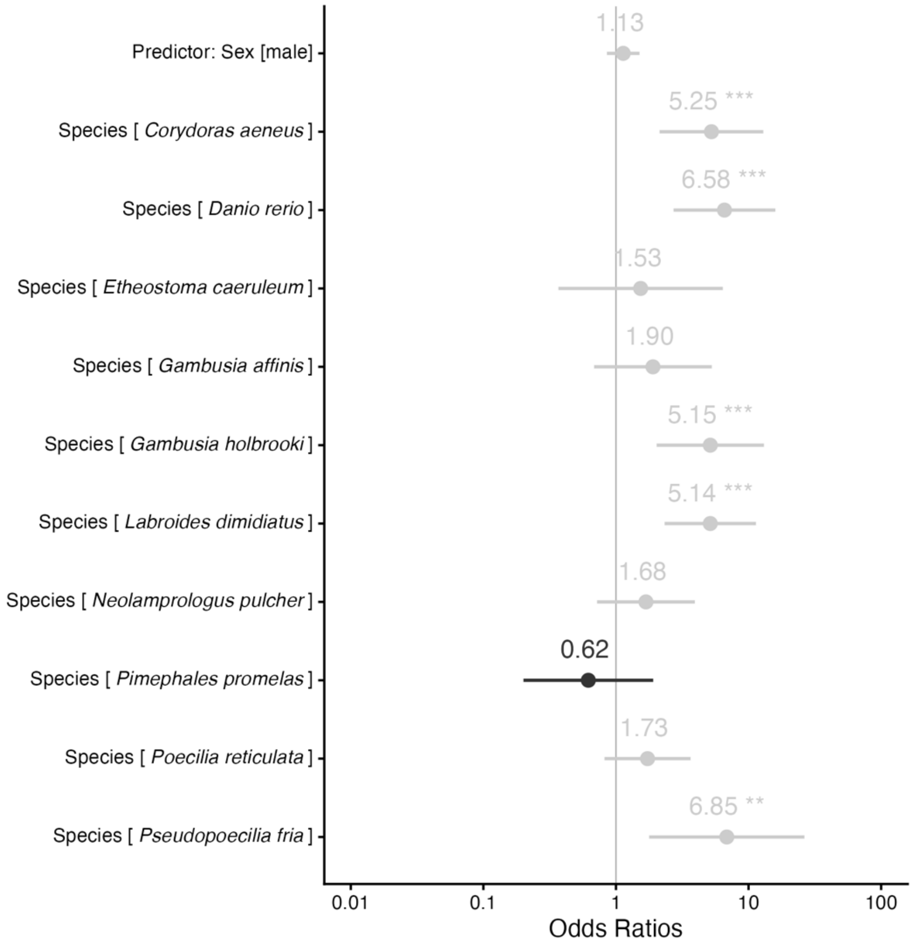


**S1 Figure.** Odds ratios obtained from the full model that included sex and species (for which data on both sexes were available). Higher values (i.e., right of the grey line) indicate better performance.

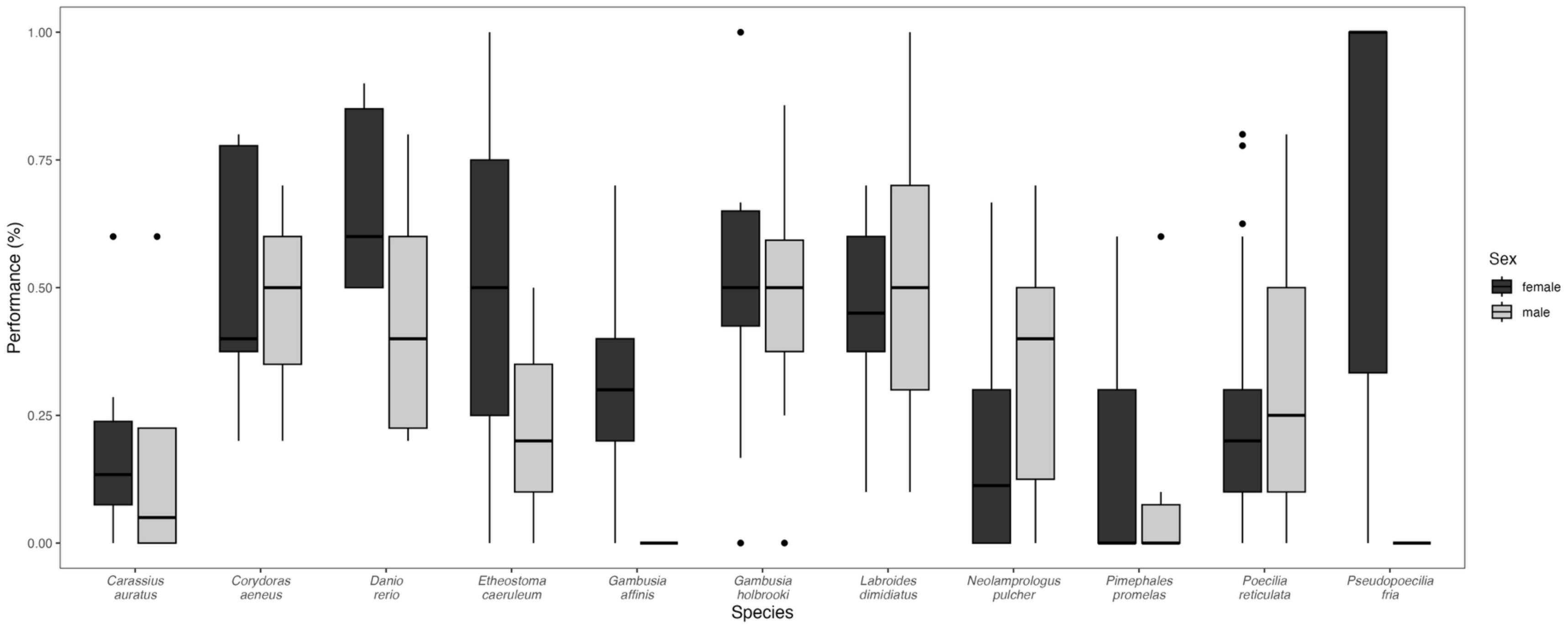


**S2 Figure.** Species' proportion of successful trials (performance) across sex. Error bars show confidence intervals. Note that only species with data on both sexes were included.

**S1 Table.** *Instruction manual for experimenters.*

| Category | Frequently Asked Question | Response |
|---|---|---|
| General setup | What size should the doors be? | The doors can be of any size (based on the aquarium dimension), as long as they allow for a full view of and easy access to the testing compartment and cylinder. |
| | What if the holding compartment varies in shape and dimension? | It is fine if the holding compartment varies in shape/dimension, as long as the testing compartment fits within the protocol requirements and this does not change the overall setup of the experiment. |
| | Can cues be used to indicate/prepare for the start of training/testing? | It can be helpful to provide subjects with light or sound cues that indicate the start of training/testing, especially when working with species/subjects that are more shy and do not readily enter the testing compartment. Note that any approach that can facilitate familiarization/conditioning with the doors and/or compartments during pre-training is appropriate. |
| Plate training (Step 1) | What is the maximum number of trials per day? | There is no maximum number of trials per day for the plate training, because it can depend on the testing site (e.g., sample size) and/or species-specific factors (e.g., food motivation, satiation levels). As a general rule, at least 6 to 10 trials per day is optimal. |
| | How much food is equivalent to a reward? | The amount of food placed on the plate + dot (or inside the cylinder during testing) during plate training will depend on the species. However, it should be roughly equivalent to one "bite", so that the fish gets the reward in only one bite (i.e., no repeated pecks). For example, if the subjects can eat 2 larvae at once, the 2 larvae can count as the "bite" unit. Although it is ideal to use as much food as possible during training (or testing), it can be difficult to use all the food during testing in some settings (e.g., facilities with strict feeding schedules). Therefore, it is fine with giving subjects the remaining daily food intake after a delay. |

| | | |
|---|---|---|
| | Where should the plate + green dot be located? | The plate should be in the same location as the cylinder during testing (for further information, see "Cylinder familiarization" section of protocol). |
| | When should the doors open? | As a general rule, the clear door should always be pulled about 2 seconds after the opening of the opaque door, regardless of the location of the fish in the holding compartment. Keeping the timing of opening consistent allows all fish species/subjects to have the same experience with the doors. |
| | When should a trial end? | A trial is considered over once the subject has eaten the reward or has not eaten it within 5 minutes. Consequently, trials in which the subject eats the food will typically be shorter than 5 minutes. Once the trial is over, the subject should go back to the holding compartment and the doors be closed (ready for a next trial if needed). Note that a trial in which the subject has not yet eaten the reward and swims back and forth between holding and testing compartments still counts as the same trial, as long as it remains within the 5-minute limit. |
| | How thick should the plate be? | It is up to the experimenter to determine the appropriate thickness (and material) of the white plate. The only requirement is that this is kept consistent throughout the steps. |
| Cylinder familiarization (Step 2) | Where should the food be placed? | During the 48 hours of cylinder familiarization, feeding can take place anywhere in the testing area (although not too close to the cylinder to avoid food getting inside). For example, the food can be placed on the right or left side of the cylinder. |
| Forced cylinder trial (Step 3) | What should the barrier frame look like? | The frame used to create the central tunnel with the cylinder can be either solid (i.e., one single piece) or made of multiple pieces, as long as it is entirely transparent. |
| | How to facilitate cylinder entrance/exit? | It is appropriate to build/add additional (transparent) structures to the aquarium if those can help with making subjects more comfortable entering/exiting the cylinder. If the subject has not passed Step 3 within at least 10 trials, opaque sides can be added. ******Addendum: If the subject has not passed Step 3 + opaque sides within at least 10 trials, it is up to the experimenter to decide to terminate the training or not (e.g., because of time constraints).* Note that during forced trials, the plate + green dot + |

| | | |
|---|---|---|
| | | food should be located in the testing compartment. This aims to motivate subjects to pass through the cylinder in order to get the food. |
| | How many times must fish enter/exit the cylinder? | Subjects must enter and exit the cylinder once. Then, they can move to Step 4. |
| Testing (Step 4) | When should a trial end? | As in Step 1, a trial is considered over once the subject has eaten the reward (whether it first touched the cylinder or not) or has not eaten it within 5 minutes. Again, once the trial is over, the subject should go back to the holding compartment and the doors be closed (ready for a next trial if needed). Note that in the case the subject enters the cylinder and eats the reward, the experimenter should allow enough time for the subject to come out of the cylinder before stopping the trial. Also, a trial in which the subject has not yet eaten the reward and swims back and forth between holding and testing compartments still counts as the same trial, as long as it remains within the 5-minute limit. **Addendum: Any deviation from the above procedure should be reported to the ManyFishes 1 Lead Team at*. |
| | How many trials per subject? | Each subject should receive a total of 10 trials. Trials in which subjects do not engage/interact with the cylinder (i.e., by not touching the cylinder and not eating the food) within the 5 minutes will still be reported but not counted towards the 10 trials. |
